# Agent-Based Modelling shows that Operation of Fisher’s Principle does not explain Zero Heritability in Human Offspring Sex Ratio

**DOI:** 10.1101/2023.04.04.535652

**Authors:** Kaitlyn T. Harper, Morgan J. Sidari, Brendan P. Zietsch

## Abstract

The evolutionary basis of the ratio of male to female births has been a focus of research for over a century. Fisher’s principle suggests that offspring sex ratios remain roughly equal due to negative frequency-dependent balancing selection: individuals with alleles that produce the rarer sex have greater fitness, increasing these alleles’ frequency until the sex ratio is balanced. However, recent analysis of Swedish births since 1932 reveals that offspring sex ratio is not heritable, challenging Fisher’s principle’s application to humans. Here, we test the hypothesis that Fisher’s principle uniquely depletes offspring sex ratio heritability at equilibrium. We used agent-based modelling to simulate Fisher’s principle over 5000 generations and found that Fisher’s principle at equilibrium does not eliminate genetic variation. These findings, along with evidence of non-heritable offspring sex ratios in humans, suggest Fisher’s principle is not a suitable explanatory framework for human offspring sex ratio.

---

Offspring sex ratio – a parent’s ratio of male to female births – has garnered interest and debate for more than a century. It is also the subject of Fisher’s principle (Fisher, 1930), which traces back to Darwin (1871) and Dusing (cited in Edwards, 1998) and is sometimes referred to as Dusing-Fisher theory. Fisher’s principle states that in sexually reproducing species, the offspring sex ratio will stabilise around 1:1 due to sexual selection favouring whichever sex is less common. This process is an example of frequency-dependent balancing selection: all else being equal, individuals genetically predisposed to producing the scarce sex will tend to have greater fitness (more grandchildren), so those individuals’ alleles will increase in frequency until the population sex ratio rebalances and the advantage of the allele disappears. According to the theory, this process keeps the sex ratio approximately even (under the condition that male and female offspring require equal parental investment).

Fisher’s principle has been the primary framework by which human offspring sex ratio has been understood (Hardy, 2002; West, 2009). Recently, though, using data from the whole Swedish population born 1932 or later (3,543,243 individuals and their 4,753,269 children), Zietsch et al. (2020) showed that offspring sex ratio is not heritable in humans. That is, genetic differences do not influence the sex ratio of individuals’ offspring, and therefore variation in offspring sex ratio cannot be inherited. Even within individuals there was no consistency at all in the sex of subsequent offspring, suggesting that offspring sex was determined by random Mendelian segregation of sex chromosomes. These observations are incompatible with the operation of Fisher’s principle – offspring sex ratio must be heritable to respond to selection as Fisher described.

If Fisher’s principle cannot operate in humans, how can it serve as a framework for understanding human sex ratio? Some have argued that Fisher’s principle makes no prediction about the heritability of offspring sex ratio at equilibrium (i.e. a stable state reached through the operation of a process) and therefore can still serve as a valid explanatory framework (Lehtonen, 2021; Orzack & Hardy, 2021). The simplest explanation under this view is that the operation of Fisher’s principle has eliminated the genetic variation in offspring sex ratio, since it must have been heritable for Fisher’s principle to operate, but is no longer heritable. While negative frequency-dependent selection is typically expected to actively maintain genetic variation (Brisson, 2018; Clarke, 1979; Zietsch et al., 2021)(Asmussen & Basnayake, 1990; Trotter & Spencer, 2007; Villanea et al., 2015), Fisher’s principle is a unique process, and it has not been established what effect the operation of Fisher’s principle has on genetic variation in offspring sex ratio. Given the complex underlying mechanisms of offspring sex ratio evolution, which are not always obvious, it is important to investigate this specific mechanism. Here, we used agent-based modelling to simulate the operation of Fisher’s principle and its effects on genetic variation and heritability in offspring sex ratio.

## Method

We created an agent-based model in R to test whether the operation of Fisher’s principle depletes genetic variation in offspring sex ratio (RCore, 2016; RStudio, 2022). The model was comprised of male and female agents (2000 total), each possessing a small genome of 20 loci which influenced their tendency to produce male or female offspring. We enabled agents to reproduce over 5000 generations, allowing us to demonstrate Fisher’s principle and observe its long-term effects at equilibrium. We decided to use 20 loci to ensure that offspring sex ratio was modelled as a quantitative trait (as opposed to a discrete trait with one or two loci). Computational load was also considered when deciding on number of loci, number of generations, and population size. In the following sections, we describe how agents were created and the rules by which they reproduced. All code and results can be accessed at https://osf.io/cbpxr/?view_only=5c437beb09534d4e8b701045e16faf01.

### Creating Initial Generation

The initial population of 2000 agents was generated, 40% were males. Each agent had 20 loci, each with two alleles, that influenced offspring sex ratio. Initial allele values were generated as either 0 or 1 (low variant or high variant), which occurred with a probability that reflected the initial population sex ratio. In the reported model, the initial population sex ratio was 40% male, so the allele values were randomised with a 0.4 probability (such that genetic scores reflected a 40% probability of producing a male offspring). Genotype was then calculated as *allele 1* + *allele 2*, thereby giving agents genotype values of 0, 1, or 2 at each locus. Genetic score for offspring sex ratio was calculated by summing genotypes for all loci. In the reported model there were 20 loci influencing offspring sex ratio, so possible genetic scores ranged between 0 and 40.

### Partner Selection

The following operations were performed five times to create five ‘mating rounds’. Agents were paired randomly, with some surplus-sex agents missing out on partnering and reproducing for that round. Mating success for the surplus sex therefore varied randomly between agents (ranging from 0-5), a result of the number of times they were randomly excluded from the mating round. Partners’ data were then linked with the agents’ data so that values for both parents were available to determine offspring attributes.

### Determining Offspring Sex

The following process emulated a genetic predisposition toward male or female offspring with additional environmental variance, such that offspring sex was determined 80% by genes and 20% by environmental factors (to simulate a largely but not completely heritable trait, like most biological traits). For each offspring, the parents’ genetic scores were averaged and transformed to a proportion of the highest theoretical genetic score (e.g. in the reported model the maximum genetic score was 40, so a score of 10 would be transformed to 0.25). This proportionate genetic score was then multiplied by 0.8, and an additional random value between 0 and 0.2 was added to represent 20% environmental influence. The resulting score was used as the probability in a random binomial outcome where 0 = female, and 1 = male; higher scores were more likely to produce male offspring. Outcome measures were recorded for the current generation at this point.

### Reproduction

Alleles were then inherited through independent assortment. That is, at each locus, an offspring inherited one allele from each parent, determined randomly. The population matrix was then truncated (i.e. all parent information removed), leaving the offspring to become the next generation, and the process described thus far was repeated. In the reported model, each simulation was run for 5000 generations.

### Outcome Variables and Analysis

We ran the model described above 50 times and averaged the simulation results for each generation. The outcome variables of interest were population sex ratio, genetic score, heritability, and genetic variance. Offspring sex ratio was recorded as the proportion of male offspring produced. Genetic score was recorded as the mean of the raw genetic scores. Given that our data was simulated, and we had complete genome information for each agent, we used simpler calculations of genetic variance and heritability than those normally used in the broader quantitative genetics literature. To calculate heritability, i.e. the proportion of phenotypic variance accounted for by genetic variance, we used the *r*^2^ of genetic score predicting offspring sex ratio (i.e. the proportion of the agent’s offspring that were male). To calculate genetic variance, we multiplied the total phenotypic variance (variance in offspring sex ratio) by the heritability, giving us the variance in offspring sex ratio that is explained by genetic scores. We used correlations to assess trends in outcome variables over generations. For ease of presentation in the main text, we have reported our model of Fisher’s principle for 1000 generations. Extended results (with 5000 generations) can be found in the Supplementary Material.

## Results

### Sex ratio and genetic scores

To determine whether Fisher’s principle operated as expected in our model, we needed to establish whether genetic scores, and consequently sex ratio at birth, changed over time. Results demonstrated that the genetic scores diverged based on agent gender, with male agents having higher scores than female agents (see Figure 1A). This result was to be expected, given that a male is more likely to have been produced by male-making alleles, and thus will have inherited more male-making alleles themselves; likewise, females will have inherited more female-making alleles. We ran a simulation where the initial genetic scores for each sex reflected this natural separation between male and female genetic scores, and all results were comparable.

**Figure 1.**
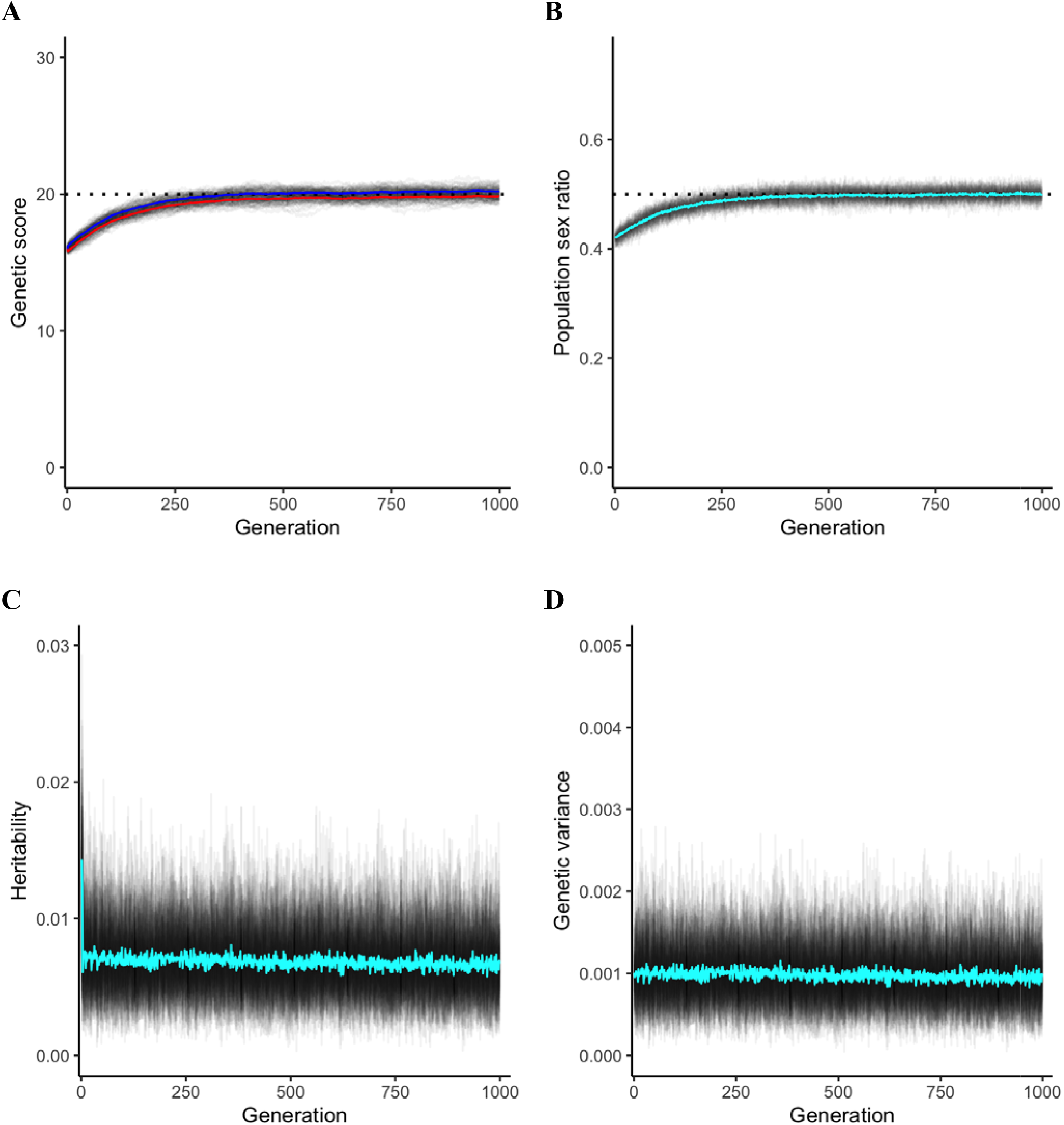
**A** Genetic score, **B** population sex ratio, **C** heritability, and **D** genetic variance over 1000 generations. *Note*. Grey lines represent individual results from 50 simulations. Coloured lines represent the mean across all simulations at each generation. In Figure B, the blue line represents the mean male genetic score and the red line represents the mean female genetic score. In Figure A and Figure B, the dotted horizontal line represents the midpoint of offspring sex ratio (1:1).

In addition to the divergence of genetic scores by sex, the mean genetic score also increased over time (from the initial mean genetic score which was biased towards female-making alleles), such that alleles for having male offspring (i.e. the rarer sex) became more prevalent (Figure 1A). The rate of this increase slowed until around generation 500 when the process reached an equilibrium at a 1:1 predisposition for offspring sex (at the midpoint of possible genetic scores – 20). The resulting population sex ratio also approached a 1:1 equilibrium (0.5 male) over the generations (see Figure 1B). The observed pattern of results thus far is precisely as Fisher’s principle would predict, where alleles leading to the less common sex increased in frequency until the sex ratio reached equilibrium at 1:1, at which point these alleles no longer conferred a benefit.

### Genetic variation and heritability

To address arguments that offspring sex ratio might no longer be heritable due to the operation of Fisher’s principle (Lehtonen, 2021; Orzack & Hardy, 2021), we next sought to test whether genetic variation and heritability were eliminated over generations.

Although heritability and genetic variance appear to be stable across generations at first glance (see Figure 1C and Figure 1D), correlations revealed that they each subtly decreased over time (*r* = -0.36, *p* <.001, for both analyses). Upon further exploration of the data, this decrease appears to be due to fixation of some alleles due to drift. An allele can fix at zero or 100% frequency due to chance fluctuations over many generations (i.e. drift), but once fixed it cannot change frequency (without mutation); that is, allelic variation can be lost but not replenished in our model, and so the occasional chance fixations will cause the genetic variation to decrease slightly over time. In analyses described in the Supplementary Material, we demonstrate that the slight decreases in heritability detected in our model of Fisher’s principle are slightly less than those observed under no selection at all (and far less than observed under directional and stabilising selection). In summary, the operation of Fisher’s principle does not deplete genetic variation, but also does not entirely buffer the loss of genetic variation due to drift.

## Discussion

In the broader field of quantitative genetics, the maintenance of genetic variation throughout various forms of selection has been extensively studied (Walsh et al., 2018). The existing literature highlights the intricate and often unpredictable nature of genetic variation under different evolutionary pressures. Our research provides a focused examination of genetic variation under a specific case of negative frequency-dependent balancing selection: Fisher’s principle. Widely accepted as the foundation for explaining the 1:1 human offspring sex ratio, Fisher’s principle has been recently contradicted by findings that offspring sex ratio is not heritable in humans (Zietsch et al., 2020). Some have argued that zero heritability at equilibrium is not ruled out by Fisher’s principle (Lehtonen, 2021; Orzack & Hardy, 2021), yet it is uncertain how the heritability necessary for the operation of Fisher’s principle could have been eliminated. With broader literature suggesting that negative frequency-dependent balancing selection is expected to maintain genetic variation (Asmussen & Basnayake, 1990; Trotter & Spencer, 2007; Villanea et al., 2015), our study aimed to determine whether Fisher’s principle could be an exceptional case where a form of negative frequency-dependent selection actively depletes genetic variation over time.

In the first generation of our model, males were the scarcer sex and the population’s alleles reflected this with a bias toward producing females. Over many generations, the sex ratio and genetic scores of the population approached equilibrium at a 1:1 offspring sex ratio. This pattern of results demonstrates Fisher’s principle as expected, such that alleles that produce the scarcer sex will increase until they are no longer beneficial (once a 1:1 population sex ratio is reached). At this point, there is no selection for these alleles, which is consistent with our findings that population sex ratio and genetic scores remained stable after this point. While our model demonstrates that Fisher’s principle is operationally sound, it does not support arguments that it could have eliminated offspring sex ratio heritability over time. Fisher’s principle, in our simulations, maintained heritability for longer than directional and stabilising selection and slightly longer than genetic drift (no selection). Previous modelling shows that negative frequency-dependent selection maintains genetic variation entirely, even counteracting drift (Villanea et al., 2015), so it is unclear whether factors of model complexity such as population size and number of alleles affected the influence of drift in our model.

Regardless, Fisher’s principle cannot account for the current observable state of no heritability in offspring sex ratio while there remains abundant heritability in virtually all variable traits that have been studied, both those under selection and no selection (Polderman et al., 2015). Given our results, Fisher’s principle would predict offspring sex ratio to be more heritable than most traits; instead, offspring sex ratio is basically unique among variable traits in being non-heritable. Additionally, anticipating the possible argument that offspring sex ratio heritability may be unobservable, we note that substantial heritability in offspring sex ratio has been detected across various species (Premoli et al., 1996; Varandas et al., 1997).

Our findings suggest that the operation of Fisher’s principle cannot be reconciled with the current non-heritable nature of offspring sex ratio, thus challenging the principle’s relevance for understanding human offspring sex ratio. We suggest that sex allocation theory in humans needs to be revised, given the foundational status of Fisher’s principle in the field, as described by West (2009, p.14):

> *‘Fisher’s theory of equal investment provides the basic null model for sex allocation theory, but it is also the foundation for all subsequent theoretical developments*.*’*

Hardy (2002, p.2, emphases added) also acknowledges that Fisher’s principle underpins extensive subsequent research:

> *‘Darwin, Düsing, Fisher, and Shaw and Mohler established the fundamental principle that members of the minority sex tend to have higher fitness than members of the majority sex. They also outlined how various ecological, demographic and genetic variables might affect the details of sex-allocation strategies by modifying both the constraints and the fitness functions*. ***Modern sex-allocation research is devoted largely to the exploration of such effects***, *which connect sex ratios to many other aspects of the biologies of many species’*

Our model is a simplified representation of the real world, and as with any model, there are limitations. Discrete generations in our model are simpler than true mating environments, and our genomes of 20 loci do not represent human genetic architecture. Nevertheless, we have no reason to expect that complexities involving linkage disequilibrium or minor allele frequencies would substantively affect our findings, especially since Fisher’s description of his principle does not implicate any of these factors. We also did not investigate factors such as group size and environmental sex ratio fluctuations. For instance, some might argue that in small groups, ongoing sex-specific mortality could lead to continuous selection and faster loss of genetic variation. However, this perspective overlooks that environmental fluctuations would not alter the genetic predisposition for offspring sex ratio in the surviving individuals. Therefore, while these changes might temporarily affect the sex ratio, the population would return to its natural equilibrium in the next generation. While differential mortality between sexes during parental care might influence selection on offspring sex ratio (Kahn et al., 2015), no specific set of circumstances has been proposed that would result in a continuous reduction of genetic variance over time.

Our findings, combined with recent evidence that offspring sex ratio is not heritable in humans (Zietsch et al., 2020), suggest that Fisher’s principle is not an appropriate explanatory framework for variation in the offspring sex ratio in humans. We anticipate various arguments that Fisher’s principle may have operated to equilibrium in humans, at which point a number of mechanisms could have depleted genetic variance in offspring sex ratio, for example an invading genetic variant coding for a 1:1 offspring sex ratio. We accept these possibilities and encourage the development of new theoretical models, but we caution the use of Fisher’s principle as an assumed starting point for new theory, given the lack of evidence for its operation in humans. Moreover, the inconsistency of offspring sex ratio within individuals (Zietsch et al.) suggests a random Mendelian process determines the offspring sex ratio in humans, although it is unclear what individual or group level evolutionary processes could have led to this outcome. Regardless, such a mechanism would be far removed from Fisher’s principle as originally proposed. We anticipate that new theoretical models will be necessary to reconstruct our understanding of sex ratio evolution in humans, and we look forward to seeing progress in this field.

