## Supplementary Material for "Agent-Based Modelling shows that Operation of Fisher’s Principle does not explain Zero Heritability in Human Offspring Sex Ratio"

### 1. Drift, stabilising, and directional selection models

The following models aim to clarify that the observed slight decrease in heritability, as presented in the main document's model, is not evidence of Fisher's principle historically influencing and then diminishing heritability in offspring sex ratio. The decrease in heritability seen in our sex ratio model is due to genetic drift causing alleles to approach fixation, as each of the allele population means move from 0.4 towards 0 or 1 over generations. Chance fluctuation in allele frequencies (i.e. drift) always occurs to some extent, even in the presence of selection. Below we have provided models of 5000-generation models of Fisher’s principle, drift (no selection), directional selection, and stabilising selection to demonstrate that the slight decreases in heritability detected in our model of Fisher’s principle are comparable to those observed under no selection at all (and far less than observed under normal selection).

#### 1.1 Drift model

The genetic drift model used the same code as the model we report in the main document with the exception that the genes in this model did not influence any outcomes. Offspring sex was randomly determined with a probability of 0.5. The traits in this model were calculated in the same way that the male offspring probability was calculated in the main model; 20% environmental variance was added to the parents’ averaged genetic scores, which were then used to assign offspring a trait value of 0 or 1. Heritability was calculated as the *r*^2^ of an individual’s genetic score predicting their offspring trait ratio (proportion of offspring with a trait value of 1), and genetic variance was calculated as the variance in phenotype (offspring trait ratio) multiplied by the heritability, giving us the variance in phenotype that is explained by genetic scores. All other processes occurred as described in the main text. This allows us to observe genetic drift occurring under no selection.

#### 1.2 Directional selection model

The directional selection model used the same code as the model we report in the main document with the exception that the focal trait is unrelated to offspring sex and is under directional selection. Offspring sex was randomly determined. The traits in this model were calculated in the same way that the male offspring probability was calculated in the main model; 20% environmental variance was added to the parents’ averaged genetic scores, which were then used to assign offspring a trait value of 0 or 1. Directional selection pressure was then applied such that 10% of the population were removed from the population before reproduction. Only agents with a trait value of 1 were eligible to be removed due to this selection. When a generation consisted of fewer than 10% of agents with a trait value of 1, all agents with a trait value of 1 were removed. All calculations were performed as described in the drift model above. This allows us to observe the depletion of genetic variation and heritability under directional selection.

#### 1.3 Stabilising selection model

The stabilising selection model used the same code as described in the directional selection model above with the exception that agents genetic scores were transformed such that scores closest to the midpoint (i.e. genetic score = 20) were more likely to produce an offspring with a trait value of 1. Thus, when the same selection pressure was applied as described in the directional model, this created a stabilising selection where middle genetic scores were more adaptive. All calculations were performed as described in the drift model above. This allows us to observe the depletion of genetic variation and heritability under stabilising selection.

#### 1.4 Results

The figures below depict the population trait scores (Figure 1), genetic scores (Figure 2), heritability (Figure 3), and genetic variance (Figure 4) for all models described above. Results demonstrate that Fisher’s principle maintains heritability and genetic variance longer than other forms of selection and is comparable to no selection. Thus, Fisher’s principle does not actively deplete heritability, though the heritability still decreases slightly over thousands of generations because of very occasional fixation of alleles whose chance fluctuations escape the balancing mechanism. Could this slight decrease explain the observed zero heritability of offspring sex ratio that we currently observe? This possibility does not make sense in the context of pervasive heritability of human traits (Polderman et al., 2015), most of which would be either under selection that would deplete heritability or under no selection at all. Given our results, Fisher’s principle would predict offspring sex ratio to be more heritable than most traits; instead, offspring sex ratio is basically unique among variable traits in being non-heritable.

**Figure 1.** Population phenotype score (trait ratio) for **A** Sex ratio model **B** Drift model, **C** Directional selection model, and **D** Stabilising selection model over 5000 generations.

| **A** | **B** |
| --- | --- |
| 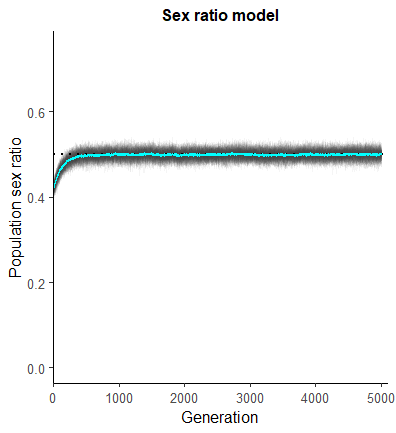  **C**  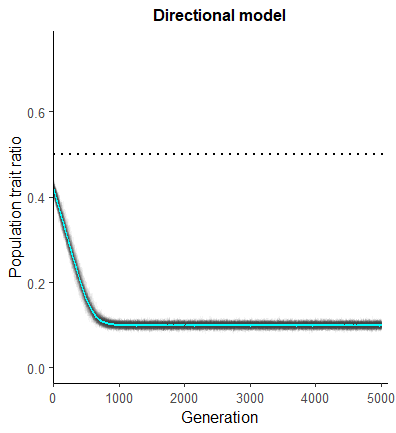 | 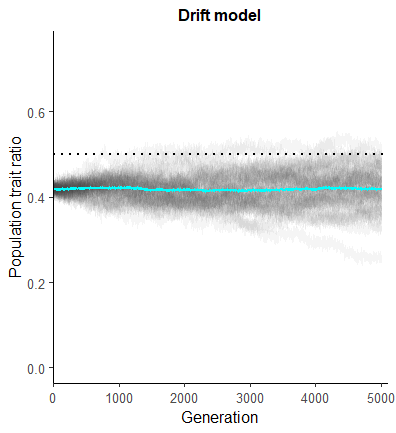  **D**  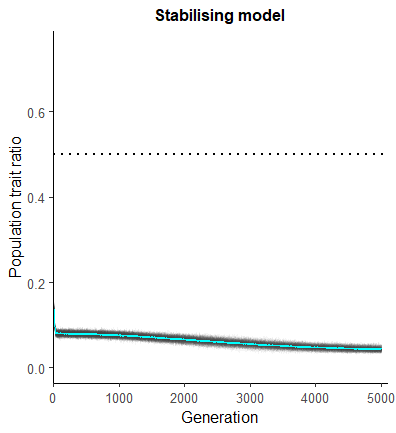 |

*Note.* Grey lines represent individual results from 50 simulations. Coloured lines represent the relevant mean across all simulations at each generation. The dotted horizontal line represents the midpoint (population trait ratio of 1:1).

**Figure 2.** Genetic score for **A** Sex ratio model **B** Drift model, **C** Directional selection model, and **D** Stabilising selection model over 5000 generations.

| **A** | **B** |
| --- | --- |
| 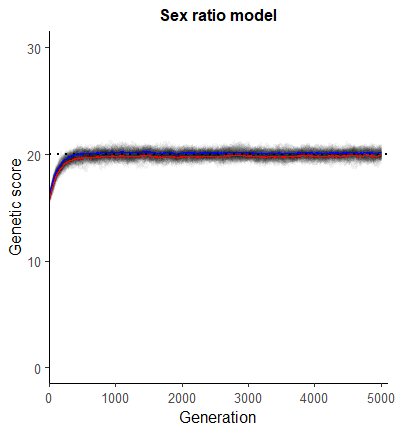  **C**  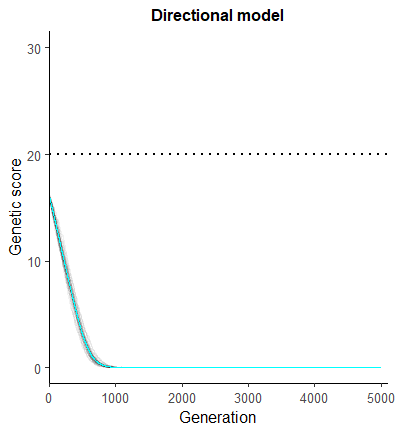 | 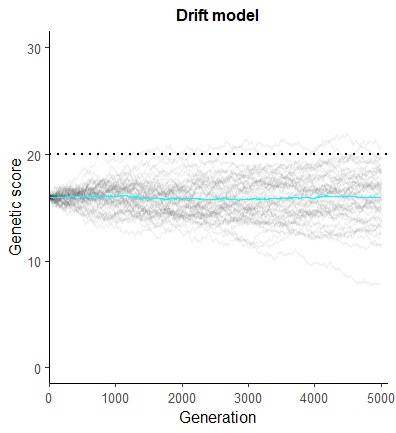  **D**  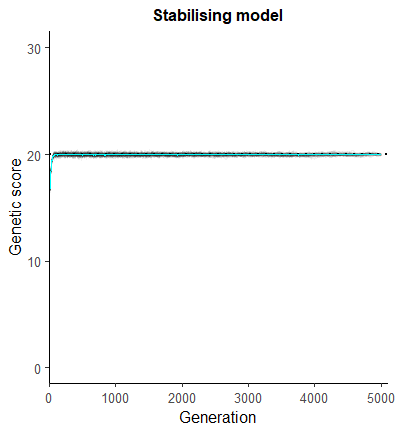 |

*Note.* Grey lines represent individual results from 50 simulations. Coloured lines represent the relevant mean across all simulations at each generation. The dotted horizontal line represents the midpoint of possible genetic scores.

**Figure 3.** Heritability for **A** Sex ratio model **B** Drift model, **C** Directional selection model, and **D** Stabilising selection model over 5000 generations.

| **A** | **B** |
| --- | --- |
| 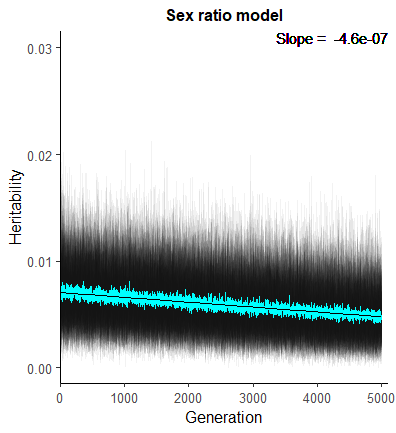  **C**  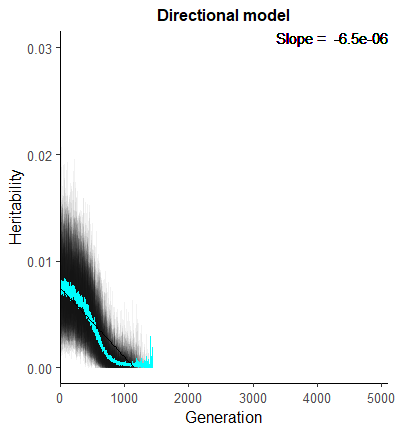 | 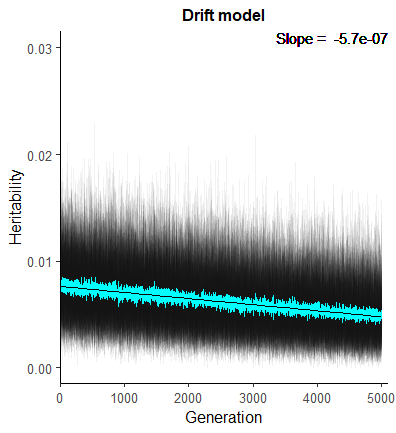  **D**  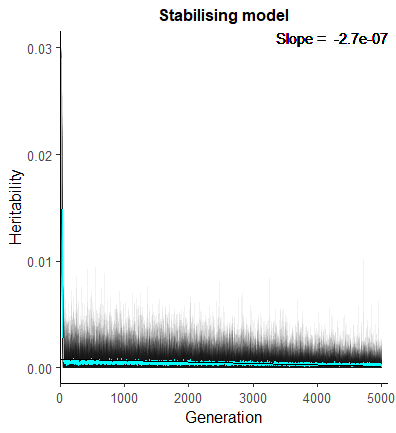 |

*Note.* Grey lines represent individual results from 50 simulations. Coloured lines represent the relevant mean across all simulations at each generation.

**Figure 4.** Genetic variance for **A** Sex ratio model **B** Drift model, **C** Directional selection model, and **D** Stabilising selection model over 5000 generations.

| **A** | **B** |
| --- | --- |
| 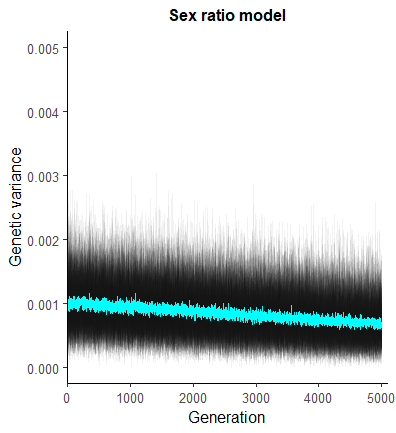  **C**  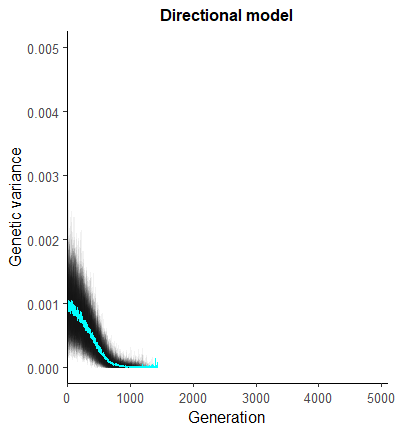 | 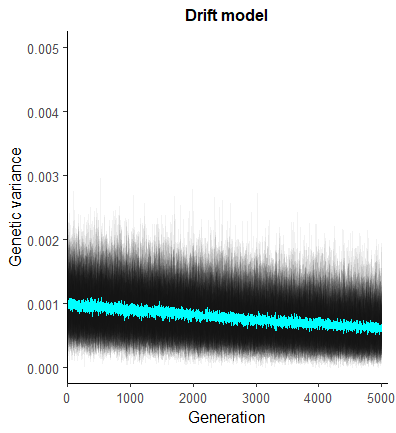  **D**  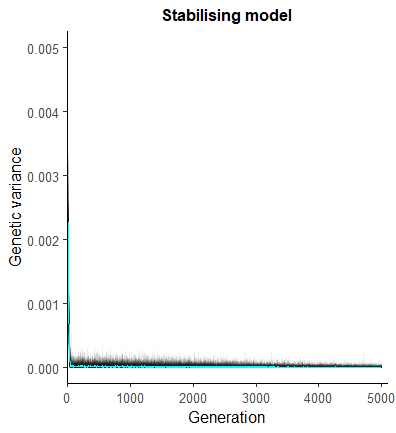 |

*Note.* Grey lines represent individual results from 50 simulations. Coloured lines represent the relevant mean across all simulations at each generation.
